# Widespread interaction between ADAR1 and transcriptional byproducts

**DOI:** 10.1101/870782

**Authors:** Chan-Shuo Wu, Sze Jing Tang, Hong Kee Tan, Li-Yuan Hung, Wei Wen Teo, Jia Li, Omer An, Kar Tong Tan, Qiling Zhou, Leilei Chen, Daniel G. Tenen, Henry Yang

## Abstract

**Background:** ADAR1, an adenosine-to-inosine (A-to-I) RNA editing enzyme, has an emerging role in cancer immunotherapy. ADAR1 presumably works by suppressing cellular innate immunity response to endogenously generated double-stranded RNAs through RNA editing. However, RNA species that are directly regulated by ADAR1 mediated RNA editing processes remain poorly defined.

**Results:** In this study, we used a novel bioinformatics approach to track ADAR1-RNA interactions. By integrating DNA-seq, RNA-seq, and ADAR1 RNA immunoprecipitation sequencing (fRIP-seq) data of K562 cell line, we provided the first *in-situ* landscape profiling of ADAR1 RNA binding and editing activities. With long RNA fragments captured by ADAR1 immunoprecipitation, we were able to identify exon junctions and genomic boundaries used by ADAR1-associated RNAs and thus we could possibly trace pre-RNA processing steps that had been acting on them. Our methodology allowed us to acquire the knowledge of transcriptome-wide scenario of ADAR1 activities. Intriguingly, we found that ADAR1 had a tendency to interact with transcriptional byproducts originated from obscure regions such as introns and intergenic regions.

**Conclusions:** Our observation might shed light on the dual role of ADAR1 proteins not only in diversifying the transcriptome, but also in reigning RNA debris from obscure regions. Moreover, as the functional potential of seemly transcriptional byproducts is just beginning to emerge, this study would bridge ADAR1 with other fields of RNA biology.

## Background

Adenosine deaminases acting on RNA (ADARs) target double-stranded RNAs (dsRNAs) and convert adenosine to inosine (A-to-I or A>I) (1). In human transcriptome, A-to-I editing is one of the most wellknown transcriptional modifications. Millions of human A-to-I editing sites have been reported by comparing RNA-seq data with the DNA reference (2-4). A-to-I editing leads to A/G mismatches (A-to-G or A>G) in the RNA-seq data and can thus be accurately identified by bioinformatics pipelines using rigorous artifact removal steps (5-7). The mammalian genome encodes three ADAR proteins, ADAR1, ADAR2, and ADAR3. ADAR1-mediated A-to-I editing is the most common type of RNA editing in mammals (5). In non-coding regions, A-to-I editing events are typically associated with the ADAR1 protein (8). Repeat elements and their surrounding regions are hotspots for A-to-I editing (2, 4, 6, 9, 10), as inversed repeat elements provide a long dsRNA structure to recruit the ADAR1 protein (5). Indeed, ADAR1 binding sites identified by cross-linking immunoprecipitation sequencing (CLIP-seq) data are mostly on ALU elements (11). ADAR1 binding is especially common for the RNAs that contain two ALU elements running in different directions (i.e., an inverted ALU pair) (12).

A-to-I editing has many possible consequences. It can modulate proteome diversity, RNA processing, RNA localization, RNA degradation, chromatin remodeling, and immune response (5, 12, 13). ADAR1-mediated RNA editing, in particular, can mark endogenous dsRNAs, avoiding them to be mistaken as foreign invaders by the innate immune system (14). Mutations in ADAR1 cause Aicardi-Goutières syndrome which can be linked to systemic autoimmune disorders (15). Protection against autoimmunity is recently recognized as the essential role of ADAR1 (5). In mouse models, the lethal phenotype of editing-deficient ADAR1 is caused by the accumulation of non-edited endogenous dsRNAs which can trigger the antiviral cellular immune response (14, 16). Hence, ADAR1 is expected to be a promising target to improve cancer immunotherapy treatments due to its regulatory role in the immune system (5, 16-18). ADAR1’s RNA targets might hold the key to fine-tuning the immune response. Currently, RNA-seq data are used for identification of RNA editing events, while detection of precise RNA binding sites of a RNA binding factor relies mainly on CLIP-seq data. As fragments in RNA-seq data are limited to a few hundred of base pairs and CLIP-seq data are concentrated on binding peak regions of <100bps, both data *per se* can only provide limited information or zoomed-in view about ADAR-bound RNAs. A broad view with deeper understanding of RNA types bound by the ADAR1 protein is thus desired.

Most of A-to-I editing sites are detected in poorly annotated and non-conserved genomic regions, such as introns, intergenic regions (IGRs), and 3’ UTRs (5, 6). Moreover, RNA editing events are co-transcriptional and rarely happen on mature mRNAs (19). Gene annotations (i.e., GENCODE, UCSC genes, or RefSeq genes) contain only mature RNA structures and thus cannot be directly used for associating editing sites with the RNA properties. To infer the properties of RNAs with ADAR1-mediated RNA editing, we employed ADAR1 fRIP-seq data to localize genomic junctions/boundaries used by these RNAs, as a gentle sonication used in fragmentation of fRIP allows long RNA fragments to be pulled down during the RNA-immunoprecipitation (20). Thus, we were able to capture the disruption of fRIP-seq signals at boundaries of defined genomic features. In this study, we established the relationship among RNA editing, ADAR1-bound RNAs, proximal genes and pre-RNA processing events by investigating whether ADAR1 fRIP-seq signals were extended or disrupted across genomic features. Our integrative analysis showed that, in CDS (coding sequence) regions, RNA-editing mediated by ADAR1 was likely discarded in mature mRNAs, while on the other hand, ADAR1 binding tended to take place in excised introns, upstream antisense RNAs, and cleaved downstream RNAs. As the functional importance of these byproduct RNAs has been recently proposed (21, 22), our findings may shed new light on ADAR1-mediated novel mechanisms.

## Results

### High concordance between ADAR1 binding regions and A-to-I editing sites

#### Identification of A-to-I editing sites in K562 cell line

We used the RNA-editing identification strategy, described in “Materials and methods”, by comparing K562 RNA-seq and DNA-seq data and successfully identified 297,407 editing sites, among which 286,393 (96.3%) were A-to-G sites while 11,014 (3.7%) were non-A-to-G sites. To further enrich ADAR1-associated editing sites, we excluded 2,208 A-to-G sites that could also be detected in ADAR1 depleted K562 cells. This resulted in a final list comprising 284,185 A-to-I editing sites for downstream analysis (Fig. S1 in Additional file 1 & Table S1 in Additional file 2). Statistical survey of these A-to-I editing sites indicated that the majority of them were in introns, intergenic regions, or 3’ UTRs (Fig. 1A & Table S1 in Additional file 2), which is consistent with the data from the popular RNA editing databases, such as RADAR and REDIportal (2, 3, 6).

**Fig. 1.**
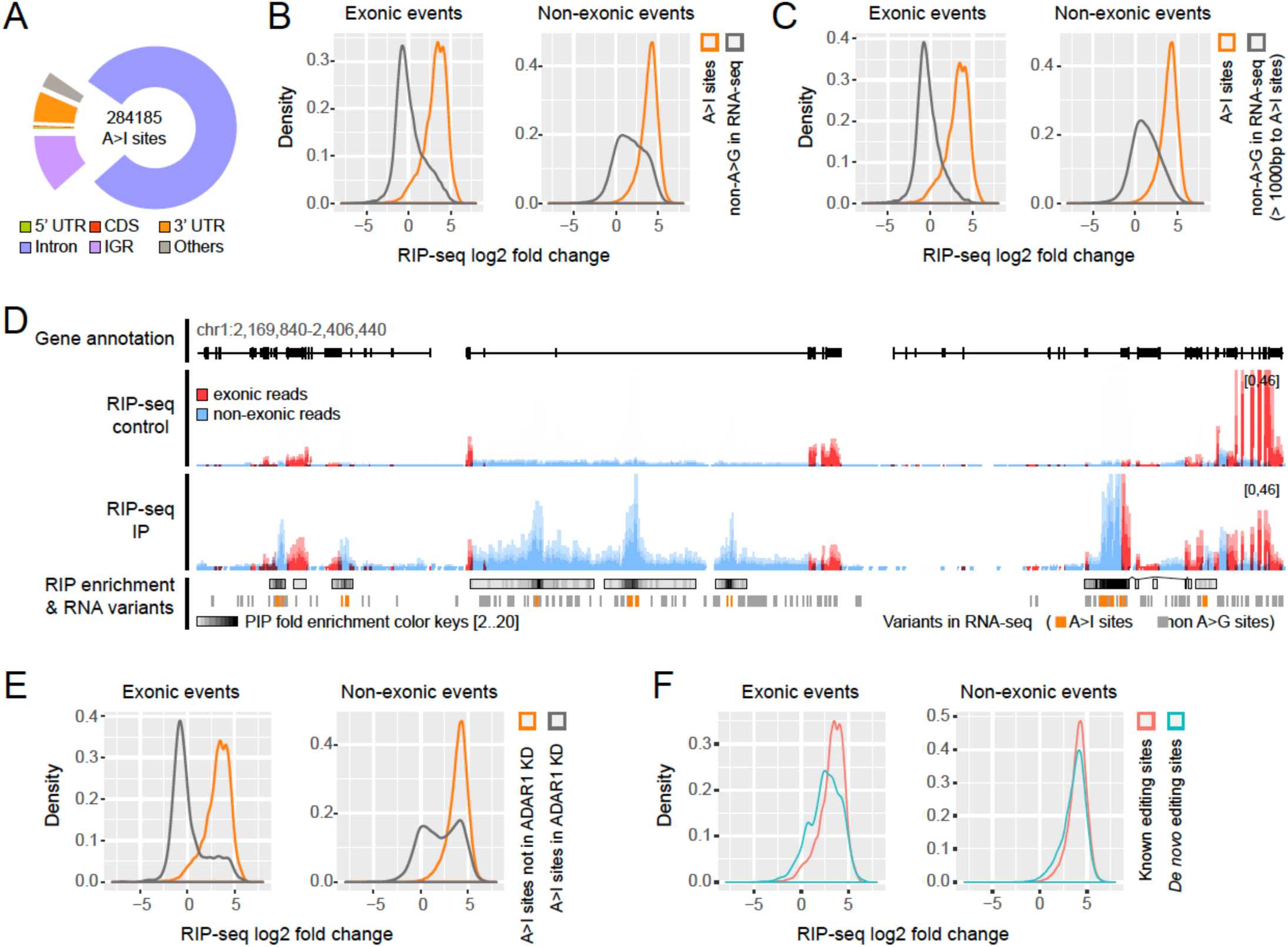
ADAR1 fRIP-seq data in K562 evaluated with RNA editing sites. (A) Genomic distribution of K562 A-to-I editing sites identified by our pipeline. List of abbreviations: UTR: untranslated region; CDS: coding sequence; IGR: intergenic region. (B) Distributions of fRIP-seq IP/control read fold changes for A-to-I editing events as well as non-A-to-G editing/SNV sites detected in RNA-seq in both exonic and non-exonic regions. In additional to non-A-to-G editing sites, RNA-seq detectable SNV sites were used to get sufficient non-A-to-I events. (C) Distributions of fRIP-seq IP/control read fold changes for non-A-to-G editing/SNVsites without neighboring A-to-I editing sites within 1k bps detected in RNA-seq along with Ato-I editing sites in both exonic and non-exonic regions. (D) Genome browser tracks of fRIP-seq data and RNA variants. Reads contained in the exonic regions (GENCODE annotation) are colored in red. Other reads are colored in blue. A-to-I editing sites are in orange. Non-A-to-G sites/SNVs are in grey. Note that non-A-to-G sites here include RNA-seq detectable SNV sites. (E) Distributions of fRIP-seq IP/control read fold changes for non-ADAR1 A-to-I editing events along with ADAR1 A-to-I editing sites in both exonic and non-exonic regions. (F) Distributions of fRIP-seq IP/control read fold changes for A-to-I editing sites supported by RADAR/REDIportal along with *de novo* A-to-I editing sites in both exonic and non-exonic regions.

#### High concordance between ADAR1 fRIP-seq data and A-to-I editing sites

ADAR1-mediated RNA editing of a site can only happen after ADAR1 has been recruited to the site (12). Thus, there should be a concordant occurrence of ADAR1-mediated RNA binding and editing. To assess such a concordance, we used the read abundance of fRIP-seq (IP & control) instead of using fRIP-seq peaks, as the RNA binding peaks could be strongly dependent upon the peak calling methods as well as the cut-off parameters applied. For each editing site, we calculated the fold change of fRIP-seq reads in the IP sample over the control. Fig. 1B depicts the distribution of such fold changes of 284,185 A-to-I editing sites and all RNA-seq detectable non-A-to-I sites in both exonic and non-exonic regions, respectively. In both regions, ADAR1 fRIP-seq reads were significantly enriched at A-to-I editing sites in the IP sample compared to the control. In contrast, most of non-A-to-I sites showed no apparent fRIP-seq enrichment, yet with viewable positive skewness (Fig. 1B). To better understand the positive skewness of fRIP-seq enrichment signals at non-A-to-I editing sites, we next selected those non-A-to-I editing sites without any neighboring A-to-I editing sites within the 1kb up- and downstream proximity, and profiled the fold enrichment distributions of fRIP reads as before. We observed no more clear skewness for those non-A-to-I editing sites without neighboring A-to-I sites in both exonic and non-exonic regions, suggesting that the positive skewness of non-A-to-I editing/SNV sites is due to the neighboring A-to-I editing sites (Fig. 1C). As long RNA fragments are pulled down in fRIP (20), the fRIP-seq signals could be extended broadly around the exact ADAR1 binding sites. As a result, some of non-A-to-G variants/editing sites at the flanking segments are also enriched or hitchhiked with the fRIP signals from their neighboring A-to-I editing sites (Fig. 1D).

Next, we checked the fRIP-seq read enrichments at those non-ADAR1-mediated A-to-I editing sites based on ADAR1 depleted RNA-seq data (Fig. S1 in Additional file 1). As A-to-I editing sites obtained from RNA-seq data of ADAR1 KD were not related to ADAR1 binding, all those editing sites were not supposed to be correlated with ADAR1 fRIP reads. Interestingly, bimodal-like distributions were observed for fRIP-seq read enrichments across those non-ADAR1 A-to-I editing sites (Fig. 1E). A-to-I editing sites in the first peak were clearly located around log_2_ fold changes of zero, thus suggesting that those sites forming the first peak were non-ADAR1 but likely ADAR2-specific editing sites. It should be noted that the ADAR1 fRIP-seq data was derived from the ADAR1 WT sample. Thus, the editing sites in the second peaks should be the common editing targets of ADAR1 and ADAR2 (Fig. 1E). Interestingly, the second peak for exonic regions was barely noticeable while both peaks in non-exonic regions were comparable, suggesting that ADAR1 and ADAR2 might share a much higher portion of common editing targets in non-exonic regions than that in exonic regions. Indeed, it has been reported that ADAR2, rather than ADAR1, is the primary editor of coding A-to-I events (8). All the above observations suggest that A-to-I editing sites in K562 are highly correlated with ADAR1-RNA binding. It is worth to note that a high percentage (∼81%) of our K562 A-to-I editing sites can be found in the RADAR/REDIportal databases. The remaining 53,075 novel A-to-I editing sites were also well supported by ADAR1 binding demonstrated with high fRIP-seq fold enrichment comparable to the sites found in these databases (Fig. 1F).

#### Determination of ADAR1-RNA binding peaks and ADAR1-mediated editing sites

Next, we applied the peak calling method (see “Materials and methods” section) for identification of ADAR1-RNA binding peaks using different IP/control fold changes as cut-off values. More than 95% of the A-to-I editing sites were covered by ADAR1 peaks with a two-fold IP/control enrichment cut-off. Even with a stringent cut-off of five-fold, the coverage of the A-to-I editing sites with fRNA-seq peaks still reached over 90% (Fig. S2 in Additional file 1). Again, these results suggest that the majority of RNA editing activities in K562 is ADAR1-mediated and can be reasonably described by fRIP-seq data. To account for most of the editing sites, we used the fold change cut-off of two for calling ADAR1 peaks in the following study. Next, we profiled the RNA binding peaks on a genome-wide scale (Table S2 in Additional file 2). The genomic distribution of peak summits (Fig. 2A) matched well with that of the K562 A-to-I editing sites (Fig. 1A). Most peak summits located in introns, intergenic regions, and 3’ UTRs (Fig. 2A). Of note, approximately 89% of ADAR1 peaks contained ALU repeat elements, in which 80% of them could form inverted ALU pair in these peaks (Fig. 2B & Table S2 in Additional file 2), indicating that ADAR1 preferentially binds to RNA regions where dsRNA structures can be formed (23, 24). This also implies the reliability of our identification of ADAR1 peaks.

**Fig. 2.**
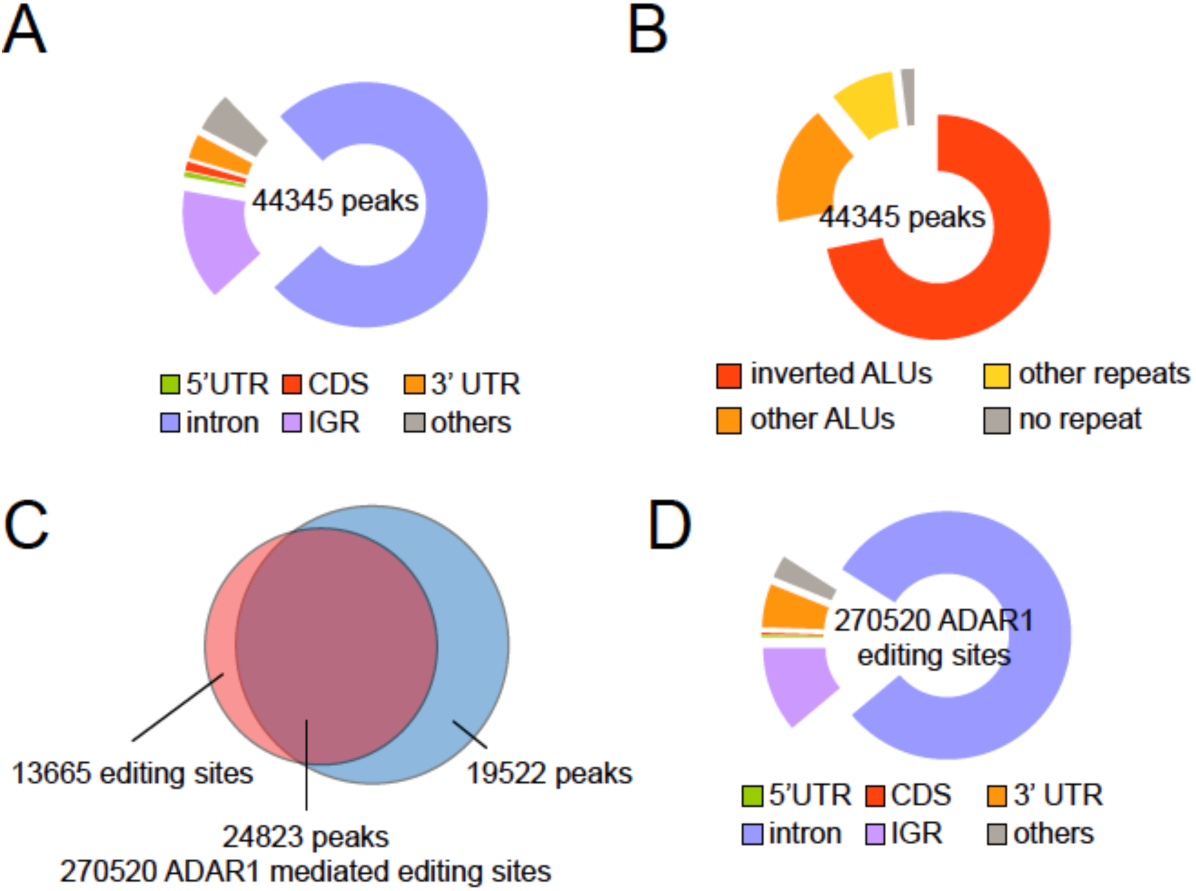
Characterization of ADAR1-RNA binding peaks in K562. (A) Genomic distribution of ADAR1 peaks. (B) Distribution of ADAR1-RNA binding peaks in terms of ALUs and other repeats. (C) Venn diagram indicating the overlap between RNA editing sites and ADAR1 peaks. (D) Genomic distribution of ADAR1-mediated editing sites.

Next, we overlapped ADAR1 peaks with RNA editing sites to derive a set of ADAR1-mediated RNA editing sites by excluding the editing sites which are not covered by the ADAR1 peaks (Fig. 2C). The overlapping of ADAR1 peaks with RNA editing sites resulted in 270,520 editing sites which we refer to as ADAR1-mediated editing sites (Table S1 in Additional file 2). These ADAR1-mediated editing sites were mainly located in introns and intergenic regions (Fig. 2D). There were 1,771 ADAR1-mediated editing sites detected outside repeated regions; however, for >99% of them, ALUs or other repeat elements could be identified within their associated ADAR1 peaks (Fig. S3 in Additional file 1). All these findings suggested that ADAR1 binds ALUs or other repeat elements to edit adenosine(s) located within the peaks, which is consistent with a previous report that RNA editing can occur a short distance away from an inverted ALU pair which recruits ADARs to exert their editing functions (25).

### Properties of ADAR1 binding in introns

#### ADAR1 binds excised introns

As the majority of ADAR1 peaks as well as ADAR1-mediated editing sites are located in introns (Fig. 2A, D), the question is then apart from premature RNAs, whether the ADAR1 protein could bind spliced-out intronic segments. To this end, we examined ADAR1 fRIP-seq signals in introns, with a particular focus on sequences surrounding exon-intron junctions. Two exemplary ADAR1 target genes, *NCBP1* and *PIP5K1C*, with intronic ADAR1 peaks were highlighted (Fig. 3A, B). A peak consisting of non-exonic reads extended to its flanking areas crossing multiple exons and introns was detected in *NCBP1* pre-mRNA transcript (Fig. 3A), demonstrating a typical example of ADAR1 binding to intronic regions of pre-mRNAs due to the existence of the inverted ALU pair. In contrast, a peak observed in *PIP5K1C* gene was located within two exon-intron boundaries (Fig. 3B), indicative of the fact that ADAR1 may mainly bind a spliced-out or excised intron fragment in this case, as there was no indication of intron retention from the control sample. We further illustrated the concept with two peaks in *GYPA* gene (Fig. 3C). One of the peaks confined within an intron, while the other peak far away from any exon-intron junctions managed to stretch widely to flanking areas (Fig. 3C). To characterize ADAR1-bound introns, we then profiled all ADAR1 peaks containing at least one A-to-I editing site which was in close proximity (<2 kb) to either 5’ or 3’ exon-intron junction sites. Moreover, to prevent fRIP-seq signals from being mangled or interfered by different binding events, we further excluded those peaks with A-to-I editing site(s) present on the opposite side of the junction within 4 kb. A clear signal disruption at the 5’ or 3’exon-intron junctions was then observed for the peaks close to 5’ or 3’ exon-intron splicing sites, respectively (Fig. 3D). ADAR1 peaks tended to shrink abruptly at the 5’ or 3’ of exon-intron junctions, despite varying distances of the closest A-to-I editing site to the junction site (Fig. 3D, E). We also noticed that the peaks located in introns were confined within the introns, implying that ADAR1 was bound to spliced-out introns. Given that ADAR1 targets unprocessed premature RNAs, a more uniform distribution of fRIP reads across the intron-exon junctions should be expected, as fragmentation or shearing of pull-down RNAs is supposed to be random and not occur exactly at exon-intron boundaries. Provided ADAR1 binds partially to the processed premature RNAs, one would expect to observe that fRIP-seq reads from the upstream exons are enriched together with the peak of the intron bound to ADAR1. Moreover, binding on partially processed premature RNAs would not be able to give the symmetrical fRIP-seq pattern between 3’ ends and 5’ ends of introns as depicted in Fig. 3. In addition, during RNA splicing, the upstream sequence is usually processed earlier than the downstream, a more intensive 5’ end interruption of fRIP signals should be expected, which was also not observed (Fig. 3D). Altogether, it is most likely that binding of ADAR1 to excised introns contributes to the observed disruption in read distributions at exon-intron junctions (Fig. 3D, E). As fRIP-seq peaks did not drop to the background level beyond the junctions (Fig. 3D, E), it indicates that ADAR1 could bind both unprocessed nascent transcript and excised introns. It should be also noted that the dramatic decrease in read abundance at the junctions was not due to PCR duplication of those reads ending at the boundaries, as duplicated fragments were already removed at the mapping step.

**Fig. 3.**
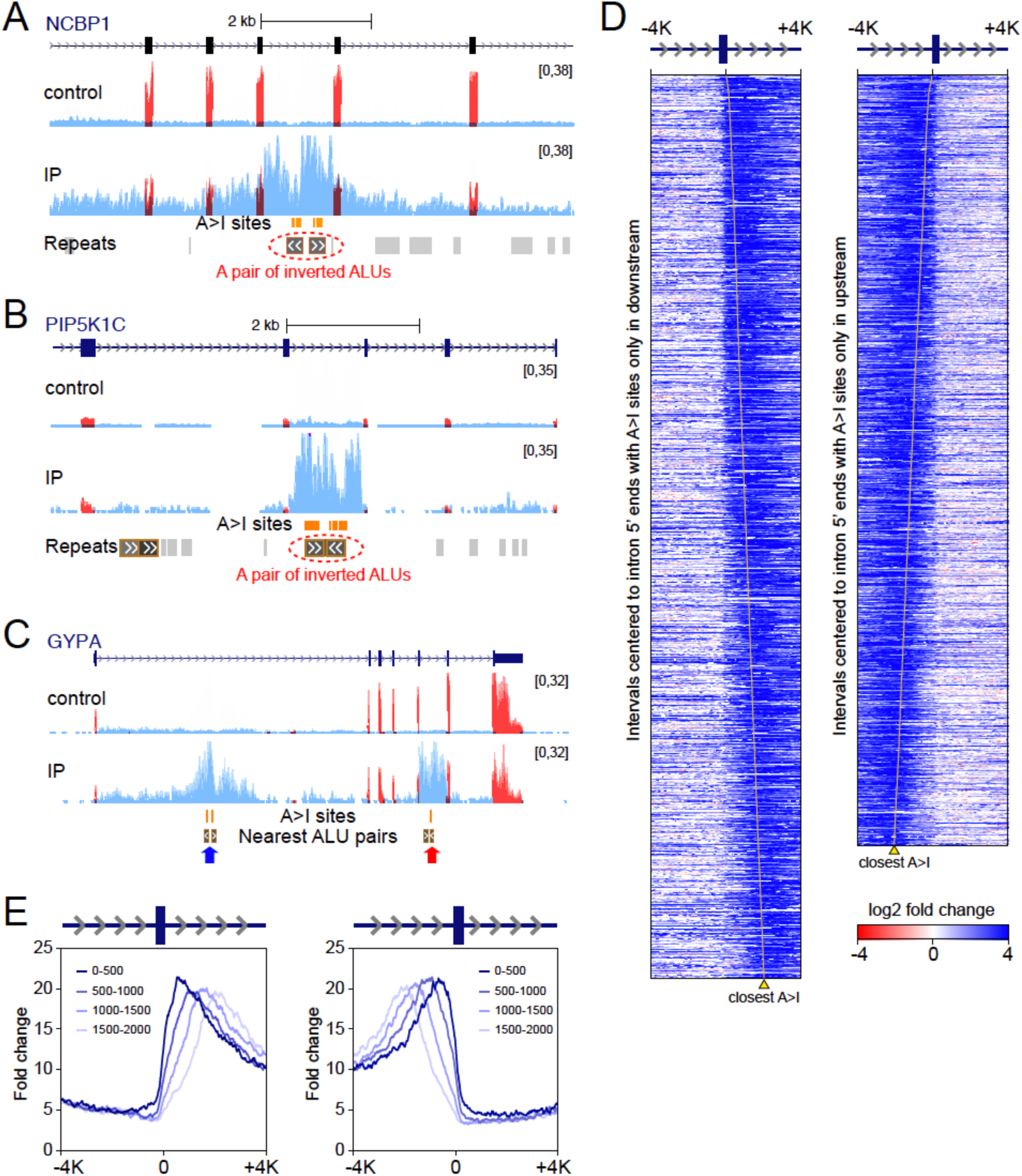
Disruption of fRIP-seq signals due to intron excision (A) Genome view of an ADAR1 peak extending to adjacent introns. The nearest ALU pair is shown for indicating the possible ADAR1 targeting locus. The track of repeats was adopted from RepeatMasker track in UCSC genome browser. (B) Genome view of an ADAR1 peak with fRIP-seq reads confined at the intron boundaries. (C) Genome view of two ADAR1 peaks. The peak indicated by the red arrow was confined within an intron, whereas the blue arrowed peak with fRIP-seq reads stretched widely to flanking areas. The nearest ALU pairs were shown for indicating the possible ADAR1 targeting loci. (D) Heatmaps of fRIP-seq read fold enrichment (IP versus control) in genomic intervals centered to exon-intron junctions for fRIP reads with at least one A-to-I site within 2,000 bps and at the same time without any A-to-I sites on the other half side of the interval. Only exons with length ≤ 2,00 bps were used. Reads were sorted based on the distances of the closest A-to-I site in the reads to the corresponding intron boundary. The yellow line indicates the position of nearest A-to-I site of each read. (E) Profiles of RIP-seq read fold enrichments (IP vs control) across intron boundaries in Fig. 3D. The profiles are divided into four groups according to the distance between the junction and the closest A-to-I site.

#### ADAR1 may not regulate splicing directly

A small proportion of RNA editing events have been reported to affect splicing regulation or vice versa (19, 26, 27). Moreover, global splicing changes have been observed after ADAR knockdown (28). To check whether ADAR1 binding is involved in splicing regulation, we examined the differential splicing events in ADAR1 KD versus the wildtype (WT) control. Indeed, differential splicing events were more abundant in the ADAR1 KD sample compared to the WT control (Fig. S4A in Additional file 1). However, only 35% of differential splicing events were overlapped with ADAR1 peaks, and the overlapping rate was comparable to the background overlapping (Fig. S4B in Additional file 1), indicating that the overlapping of splicing events with ADAR1 peaks might be random and not specifically linked to a direct regulation of ADAR1 on splicing. Further, by conducting the gene ontology analysis for differentially spliced genes between the ADAR1 KD sample and the WT control. “Spliceosome” was the most significantly enriched pathway (Fig. S4C in Additional file 1), suggesting that ADAR1 may affect expressions of spliceosome-related genes, which might eventually lead to the aberrant splicing. In sum, it might be less common for ADAR1 to directly regulate RNA splicing; instead, ADAR1 may mainly regulate splicing through regulating a subset of spliceosome-related genes.

### Properties of ADAR1 binding in exonic regions

#### ADAR1 binds mature RNAs at 3’ UTRs

Next we shifted our focus to RNA editing activities on exons. Well-studied and renowned editing sites are usually in exons which require the adjacent intron sequence to form dsRNA for ADAR1 recruitment (29, 30). In such cases, editing must precede splicing. In contrast, exons recruiting ADAR1 without the presence of an adjacent intron have rarely been documented. To identify such mature ADAR1-bound RNAs, we searched for ADAR1 peaks across multiple exons with exon-exon junction-spanning reads. We found 407 cases of such junction-spanning peaks. Among them, 283 (69.5%) peaks contained A-to-I editing sites with 234 ones containing inverted ALU pairs (Table S2 in Additional file 2). As expected, nearly all (96.6%) of the associated A-to-I editing sites were either in 3’ UTRs or non-coding transcripts. The majority (97.3%) of the exon-exon junction-spanning peaks contained the last two exons and bridged over the last introns. As an example, fRIP reads without introns but with both inverted ALU pair and A-to-I editing site(s) were detected at the 3’ UTR of *RAB2B* gene (Fig. 4A).

**Fig. 4.**
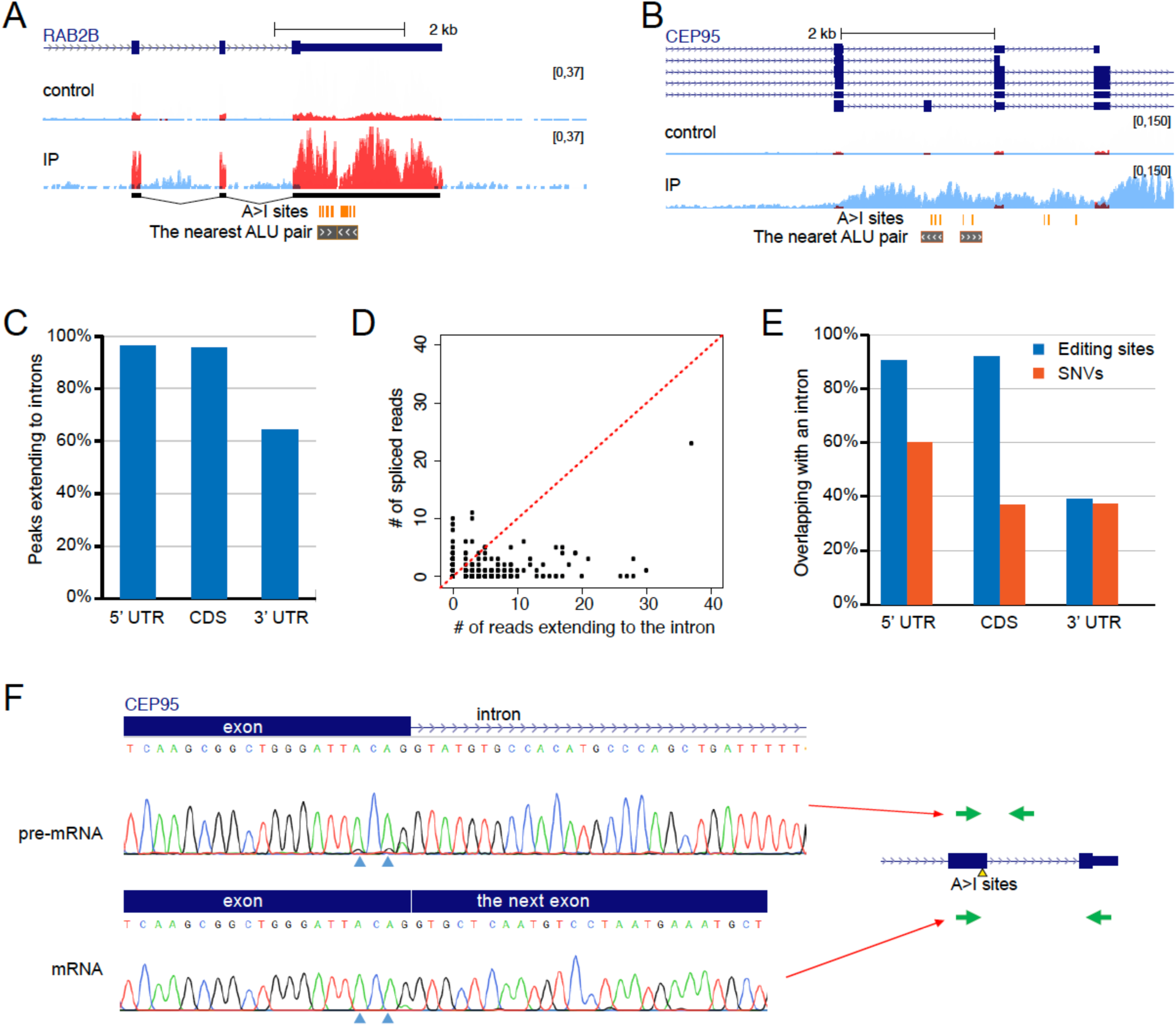
Characteristics of ADAR1 binding in exons. (A) An example of an ADAR1 peak across multiple exons. The nearest ALU pair was shown in 3’ UTR highlighting the potential ADAR1 targeting locus. (B) An example of ADAR1 binding to a GENCODE annotated exon. The nearest ALU pair is shown to highlight the potential ADAR1 targeting locus. (C) Percentages of peaks in various exonic editing sites with fRIP reads extending to introns. (D) Scatter plot of the number of RNA-seq reads with exon-exon spanning versus the number of RNA-seq reads bridging the exon-intron junction. Dots represent A-to-I editing sites located in exons and all RNA-seq considered in the plot contained the A-to-I editing. (E) Percentages of identified exonic editing sites overlapping with an intron of other isoforms in different genomic features. Exonic SNV sites are included as background for non-editing events. (F) Two A-to-I editing sites were observed in the non-spliced RNA, but absent in the mRNA. The green arrows indicate the concept of primer design.

#### ADAR1 A-to-I editing sites in coding regions may not be genuine

When an exon requires an adjacent intron to form dsRNA for ADAR1 recruitment, the corresponding ADAR1 fRIP-seq peak will extend to the intron without enrichment of exon-exon junction-spanning reads (Fig. 4B). Surveying all the fRIP peaks with exonic editing sites, we found in fact that almost all of them in 5’ UTR and CDS regions extended reads into intronic regions (Fig. 4C). As the boundaries of 5’ UTR regions could be more complicated in the annotation, we concentrated our effects on the CDS editing sites. As the supporting ADAR1 peaks with CDS editing sites did not extend the fRIP reads to exon-exon junction but mostly to neighboring introns, it was questionable whether the associated editing site(s) carried over to the mature RNAs. When surveying those RNA-seq reads possessing CDS A-to-I editing event(s) and spanning a splicing junction, we found that they tended to extend into the intron instead of spanning the exon-exon junctions (Fig. 4D), indicating that inosines tends to occur in unprocessed nascent transcriptions or excised introns. Next, we checked whether introns of other isoforms could cover those GENCODE annotated “CDS” editing sites. Truly enough, most (>90%) of these editing sites could also be located in the introns of other isoforms (Fig. 4E). Such a case was illustrated in Fig. 4B, and experimental validation of such a case was shown in Fig. 4F based on the Sanger sequencing results. When using a primer pair across an exon-intron junction targeting pre-mRNA or excised intron, two editing events could be observed. However, using a primer pair across an exon-exon junction targeting the mature mRNA, we were unable to recover the two editing events (Fig. 4F). Moreover, the ability of ADAR1 to bind excised introns provides an extra alternative explanation to these inferred “CDS” editing sites, further reducing their reliability. Hence, ADAR1 editing sites in CDS might not be necessarily associated with valid CDS editing sites, and false CDS editing sites could commonly occur from assigning them a wrong identity in the gene annotation. Careful experimental validation is thus needed when studying CDS editing sites.

### Properties of ADAR1-mediated RNAs in intergenic regions (IGRs)

#### Characteristics of ADAR1-mediated RNAs from IGRs

In addition to intronic and exonic ADAR1-mediated editing sites, we identified 30,019 ADAR1-mediated editing sites in IGRs. These editing sites were associated with 3,191 ADAR1 peaks, in which 1,229 (38.5%) of them had their fRIP signals reaching gene bodies within 500bps. The distribution of these peaks showed that ADAR1 binding sites in IGRs were more frequently found in downstream of genes than in upstream (Fig. 5A & Table S3 in Additional file 2).

**Fig. 5.**
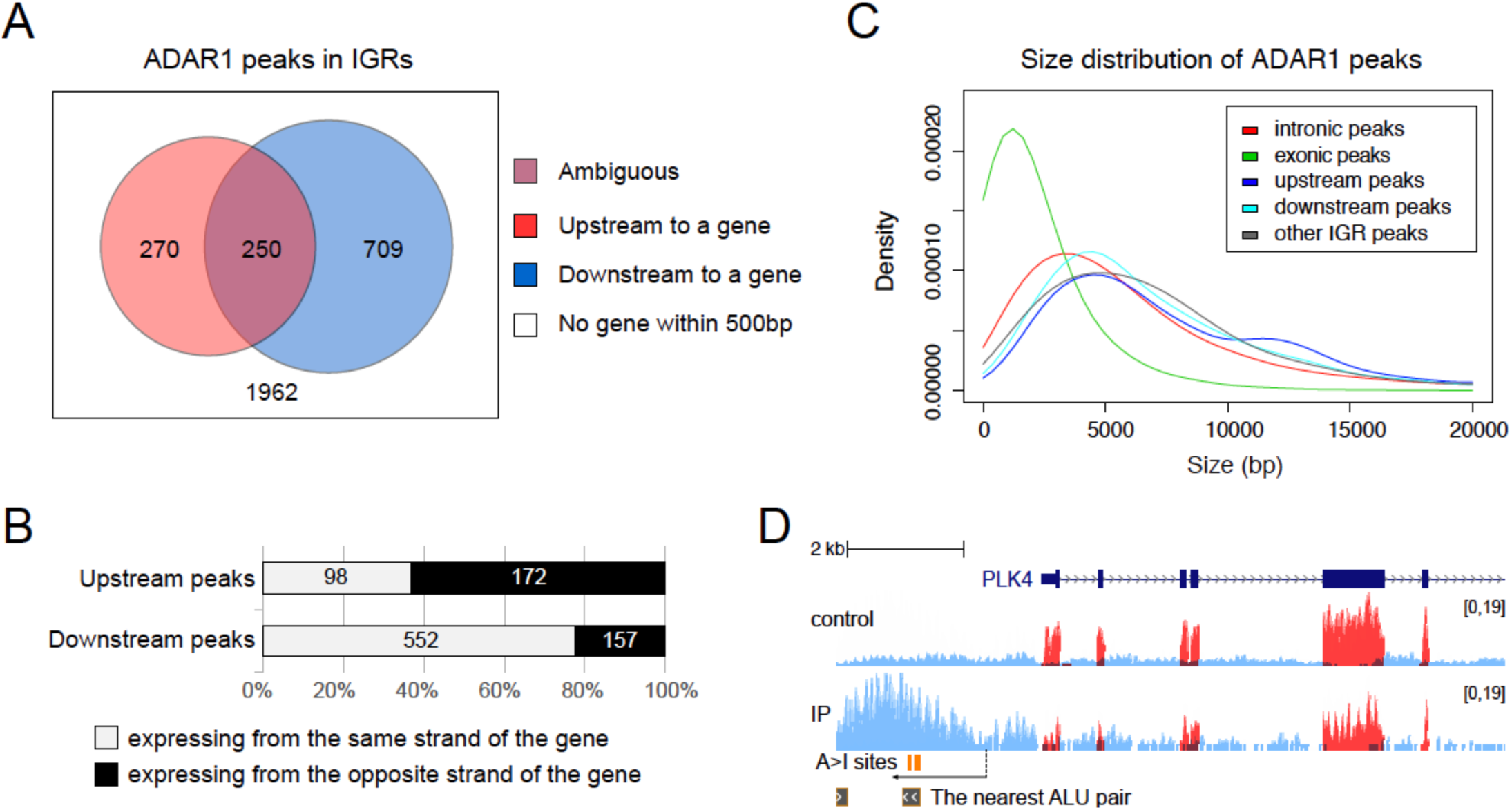
RNA binding peaks in IGRs (A) Venn diagram depicting the localization distribution of intergenic ADAR1 peaks with respective to neighboring genes. (B) Bar charts describing the expressing strand of fRIP-seq reads in upstream or downstream peaks with respective to neighboring genes. (C) Density plot depicting the size distributions of ADAR1 peaks in different genomic features. (D) An example of an ADAR1 upstream peak to a gene (*PLK4*). The arrow below A-to-I sites indicates their direction.

As the K562 fRIP-seq data were strand-specific RNA-seq, we were able to find that the reads in ADAR1 peaks upstream of genes tended to be expressed from the opposite strand of the nearby genes, while the reads in ADAR1 peaks downstream of genes were usually expressed from the same strand of their upstream genes (Fig. 5B & Table S3 in Additional file 2). Inverted ALUs were usually present (∼82%) in the IGR peaks, and those ADAR1-mediated fRIP peaks were larger compared to exonic and intronic peaks (Fig. 5C). An example of ADAR1-mediated upstream RNAs illustrates this concept (Fig. 5D).

#### ADAR1 binds cleaved downstream debris RNAs

The strong strand bias of ADAR1-mediated downstream RNAs indicates that they may closely be associated with their upstream genes (Fig. 5B). Thus, we examined those downstream ADAR1 peaks with at least one A-to-I editing site located within 2 kb downstream of an annotated polyA site on the same strand. Surprisingly, the fRIP-seq signals of these ADAR1 peaks started to be enriched right downstream of the annotated polyA sites (Fig. 6A, B). As the fold changes were used in the heatmap (Fig. 6A) and the read numbers downstream of 3’ UTR were low in the control, we asked whether what we observed was simply due to division by a small number. Thus, we replaced the fold changes with the differences of fRIP-seq read coverage between the IP and control samples, and a similar distribution also occurred exactly at the end of the polyA sites (Fig. 6C). The question is then whether the fRIP reads downstream of a polyA site belong to the same transcript or an independent one. Provided the latter were true, the 3’ UTR end would be exactly a TSS site of another independent transcript, which is unlikely the case. Hence, the fRIP reads downstream of polyA sites are most likely to belong to the cleaved fragment of the same transcript, suggesting ADAR1 may bind the RNA debris downstream of a transcript. Moreover, since the ADAR1 peaks can extend far towards the downstream, ADAR1-bound downstream RNAs can be surprisingly large (Fig. 5C, 6B & 6D). Our findings also implied that a fraction of downstream editing sites could be wrongly recognized as 3’ UTR editing sites, as an ADAR1-bound RNA mapped to 3’ UTR regions could be the downstream RNA from another isoform with a shorter 3’ UTR. An example is shown in *MRPS33* locus (Fig. 6D).

**Fig. 6.**
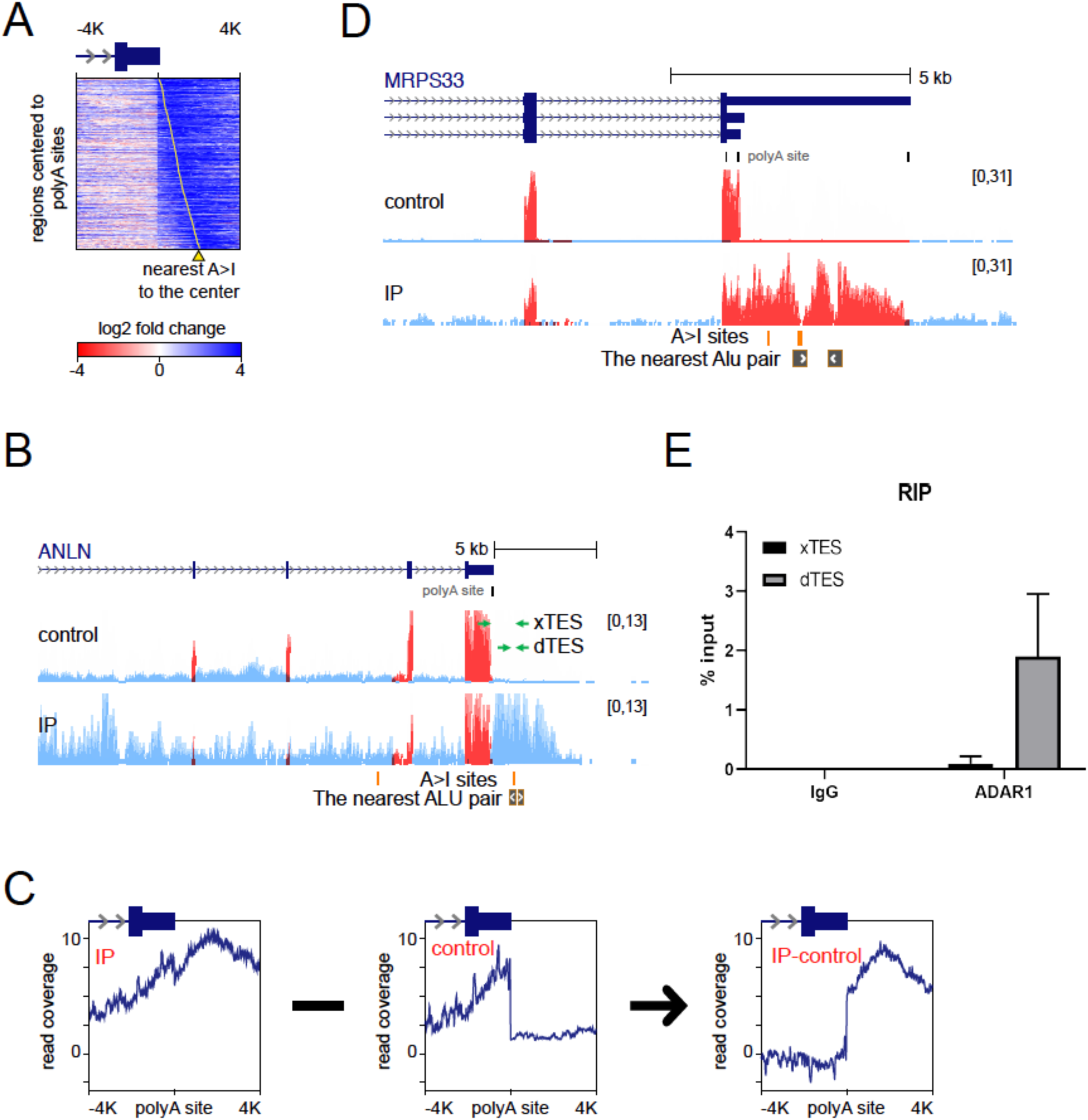
Characteristics of fRIP-seq reads around polyA sites. (A) Heatmap of fRIP-seq fold enrichments in intervals centered to transcription end sites (TESs). Only TESs annotated with polyA sites were used. The polyA site must be with A-to-I sites on the same strand within 2000bp downstream and find no A-to-I site with 4000bp upstream. (B) Genome view of ADAR1 binding at the downstream RNAs of a gene (*ANLN*). The nearest ALU pair is shown for highlighting the potential ADAR1 targeting locus. Green arrows indicate the concept of primer design for Fig. 6E. (C) Profiles of fRIP-seq read coverages in IP and control and the difference between them around TESs in Fig. 5A. The difference is the subtraction result between IP and control based the normalized read coverage. (D) Genome view of fRIP-seq around the TES of gene *MRPS33*. The ADAR1 bound downstream RNAs are annotated by GENCODE as 3’ UTR, but the ADAR1 peak was stopped at the upstream polyA site. The nearest ALU pair is shown for highlighting the potential ADAR1 targeting locus. (E) Validation of ADAR1 binding on *ANLN* locus by qPCR of ADAR1 RIP pulldown.

Next, we proceeded to experimental validation. Firstly, we performed RNA immunoprecipation with ADAR1 antibody in K562 cells. Sanger sequencing of ADAR1 pulled-down RNAs confirmed the A-to-I editing sites identified with RNA-seq data (Fig. S5 in Additional file 1). With the pair of primers across the 3’ UTR end, no visible qPCR product was found in the ADAR1 RNA pulldown, while strong and clear qPCR product was detected for the primer pair located only in the downstream RNAs (Fig. 6E). Overall, our results suggest that ADAR1 can target the downstream RNAs of transcripts, which are already disconnected from the transcripts by the 3’ UTR cleavage event.

## Discussion

Although identification of A-to-I editing sites from RNA-seq data has become a routine endeavor (6), the functions of most reported editing sites still remain largely unknown (12) as A-to-I editing sites are predominantly located in poorly annotated and non-conserved genomic regions, such as introns or intergenic regions (2, 6). This imposes difficulties on connecting them with established knowledge. In this study, we developed a methodology to study host RNAs of A-to-I editing events, and explored their relationships with genomic features of adjacent genes. Furthermore, we inferred how host RNAs of A-to-I editing events were processed during ADAR1 binding by integrating both RNA-seq and fRIP-seq data.

Firstly, we found a high concordance between A-to-I editing sites and ADAR1 RNA-binding peaks with high presence of inverted ALUs. We showed the concordance from different viewpoints by checking the distributions of fRIP-seq signals for all A-to-I editing sites versus those for non-A-to-I editing sites or SNV sites in both exonic and non-exonic regions. Reliable detection of ADAR1 peaks with fRIP-seq data was demonstrated from different angles.

Comparing RNA-seq data of ADAR1 depleted cells with ADAR1 WT ones, we were able to identify a set of ADAR2 specific A-to-I editing sites as well as ADAR1/2 common editing sites. Interestingly, the common editing sites between ADAR1 and ADAR2 in exonic regions are rare while the number of such common sites in non-exonic regions is comparable to that of ADAR2-specific editing sites. All these findings are in a good agreement with the reports that ADAR2 is the major enzyme performing RNA editing in coding regions (8), while ADAR1 is mainly for editing in the non-coding regions (8). Indeed, we also found that CDS editing sites covered by ADAR1 peaks were less reliable. They could actually be intronic editing sites overlapping with the coding regions of other isoforms. Rigorous experimental validation is thus required when studying A-to-I editing sites in coding regions.

Precise RNA binding sites of ADAR1 have been detected using CLIP-seq data and found mostly on ALU elements (11). In this study, we aimed mainly to characterize the RNAs hosting the ADAR1-RNA binding. Long RNA fragments in fRIP-seq data provide a unique opportunity to determine the identities of ADAR1-bound RNAs being premature, mature, or spliced-out by profiling fRIP-seq reads at the boundaries of genomic features such as exon-exon or exon-intron junctions, TSS or TES ends. In this study, ADAR1-mediated RNA editing sites could even be successfully associated with RNA species that were absent in gene annotation databases.

It has been reported that A-to-I editing events predominantly happen on nascent transcripts (19) and RNA editing events may affect splicing regulation or vice versa (19, 26, 27). In this study, we demonstrated that ADAR1 is likely involved in splicing indirectly via regulating splicesome genes. Thus, RNA editing and splicing events might not be directly related. We further investigated whether the RNAs bound to ADAR1 were subjected to certain pre-RNA processing steps during A-to-I editing. The pattern of fRIP-seq signals showed that pre-RNA processing could be coupled with A-to-I editing events in 3’ UTRs, introns, and even intergenic regions. Some ADAR1 fRIP-seq peaks from 3’ UTR editing events could span multiple exons, indicating that the last introns of their host RNAs were spliced out. In contrast, we did not observe any intronic editing events to extend the reads of fRIP-seq peaks to upstream exons. Instead, their fRIP-seq signals were generally disrupted around intron boundaries when they extended to upstream or to downstream. The above patterns at exon-exon and exon-intron junctions suggest that the binding of ADAR1 does not cease after the intron(s) has been spliced out, no matter whether ADAR1 binds 3’ UTRs or introns. A recent study demonstrates that excised linear introns can play a regulatory role in yeasts (22). If excised introns also have regulatory functions in human cells or ADAR1 also regulates function-rich 3’ UTRs, what kind of role ADAR1 plays in excised introns or matured 3’ UTRs would be very interesting to elucidate besides binding to dsRNAs.

ADAR1 fRIP-seq peaks guided us to associate nearly half of intergenic A-to-I editing sites with proximal genes. We found that A-to-I editing sites at the upstream of genes usually associate with long antisense RNAs. Upstream antisense RNAs share the bidirectional promoter with the downstream gene are common byproducts during gene transcription, and their functional potential is beginning to emerge (21). More interesting discoveries came with ADAR1-mediated A-to-I editing sites after polyA sites. The pattern of fRIP-seq signals at the polyA sites suggests that the corresponding editing events can happen on cleaved downstream RNAs. Again, ADAR1 interacted with a transcriptional byproduct, this time at the downstream of a gene.

Surprisingly, the interactions between ADAR1 and transcriptional byproducts seem to be widespread. It is mysterious why ADAR1 spends much of its efforts to target RNA species which are subjected to degradation. As ADAR1 peaks are relatively broad in intergenic regions and RNA editing sites are rarely fully edited, there could be a hint that ADAR1 might target a subset of transcriptional byproducts escaped from earlier RNA degradation. As long unedited endogenous dsRNAs could trigger viral immune response, ADAR1 has become an emerging target of cancer immunotherapy (16, 17). Thus, this type of transcriptional byproducts may be of clinical values and should not be overlooked. They might be the major targets of the cancer immunotherapy sensitizer, ADAR1. Understanding the interplay between them and the innate immune system would be an important future work. Our work provides important clues to study the interplay between ADAR1 and these seemingly byproducts of transcription.

## Conclusions

Integrating ADAR1 RIP-seq with long RNA fragments and other deep sequencing data, we provided the first transcriptome-wide profiling of ADAR1 activities and characterized ADAR1-bound RNAs in a systematic way. We reported the widespread interaction between ADAR1 and transcriptional byproducts originated from intronic or intergenic regions. This explains why a large proportion of A-to-I editing sites in widely-used RNA editing databases was in non-annotated genomic regions. Our observation might shed light on the role of the ADAR1 protein in reigning RNA debris from obscure regions and bridge ADAR1 with other fields of RNA biology.

## Materials and methods

### Analysis of ADAR1 binding sites in K562 cells

The ADAR1 K562 fRIP-seq data set and corresponding input control were downloaded from ENCODE (31) (Table S4 in Additional file 2), and mapped using STAR against human GRCh38 with GENCODE transcriptome annotation (32, 33). PCR duplications were removed using samtools rmdup (34). The resulting alignments for each data set were separated into two bam files. One bam files contained reads in the STAR transcriptome mapping (i.e., reads being mapped into ENCODE transcripts). The other bam file contained the other reads. Bam files were further converted to bedGraph format using the bedtools (35). Prior to the bedGraph conversion, normalization of different samples was performed based on the number of reads in the STAR’s transcriptome mapping. The larger data set (i.e., the control) was shrunk down to fit the size of the smaller one (i.e., the RIP). Comparison of the IP and the control was performed in a moving window of ±150 bp for each site. Those subsequent sites with average fold enrichment of ≥ 2 and Poisson distribution p-value ≤ 10^−5^ were summarized together as a peak for ADAR1 binding. Each peak was extended to flanking areas until the fold enrichment dropped below 2. It should be noted that a peak from the transcriptome mapping part was able to span splicing junctions. The summit of a peak was defined as the position with highest fold enrichment. The resulting ADAR1 peaks were used to represent ADAR1 binding sites in this study.

### Identification of A-to-I editing sites in K562 cells

The following ENCODE sequencing data sets from K562 cells were downloaded: strand-specific total RNA-seq, ChIP-seq input used as DNA-seq, and total RNA-seq of ADAR1-depleted K562 cells (Table S5 in Additional file 2) (31). For each data set, reads were mapped using STAR against human GRCh38 with GENCODE transcriptome annotation (32, 33). Two rounds of iterative mapping were performed. In the first round, we followed the ENCODE options in the STAR manual except the number of mismatches was limited to 5% of the read length. To account for more frequently edited regions, unmapped reads from the first round were re-mapped. At the second round, the STAR aligner was allowed to search for up to 500 anchor loci in the reference genome, and the limitation of the mismatch number was loosening to 10% of the read length. The mapping results of the two rounds were merged together. RCR duplications were removed by using Picard Toolkit (36). We further removed those multiple mapped reads, reads spanning over nonannotated splicing junctions, and reads not mapped in pairs. To call the editing sites, we followed the filtering approach described in (7) with following adjustments. We used 5% as the editing level cut-off in each data set. Candidate editing sites supported only by the leading 20 bases or the ending 20 bases of reads were removed. The ChIP-seq input files from 47 ChIP-seq data sets was used as the reference DNA-seq to remove potential SNV sites or artifacts from candidate editing sites. Furthermore, total RNA-seq data of ADAR1-depleted cells were used to reduce non-ADAR1 editing sites (Fig. S1 in Additional file 1). All results were formatted to bedGraph or bed files.

### Data visualization

The bedGraph and bed files were further converted to bigWig and bigBed using bedtools, respectively (37). The resulting files were then imported into the UCSC Genome Browser for visualization (38-40). Furthermore, reads that mapped into GENCODE transcripts were colored red (i.e., exonic reads were red), and the other reads were colored light blue (i.e., non-exonic reads were light blue). The deepTools were used to analyze bigWig files and bed files for generating signal profiles and heat maps (41).

### Native RNA immunoprecipitation (RIP) and RT-qPCR

Ten million K562 cells were lysed with 1 ml native lysis buffer [150 mM KCl, 25 mM Tris pH7.5, 5 mM EDTA and 0.5% NP-40 supplemented with 1U/μl SUPERase-In (Invitrogen) and 1x cOmplete protease inhibitor (Roche)] followed by preclearing with 50 μl Dynabeads G (Invitrogen) at 4°C for 4 hours. After pre-clearing, 5% of lysate was kept as input. Subsequently, 5μg rabbit IgG (Cell Signaling) or anti-ADAR1 antibody (Sigma, HPA003890) was added to pre-cleared cell lysate followed by overnight incubation at 4°C with rotation. Afterward, 50 μl Dynabeads G (Invitrogen) was added to mixture followed by incubation at 4°C with rotation for 2 hours. After incubation, beads were washed with native lysis buffer three times followed by RNA extraction using RNeasy mini kit (Qiagen). 12.5μl RNA was used for cDNA synthesis with Advantage RT-PCR kit (Clontech). Detection of pull-down product was performed with qPCR using GoTaq® qPCR Master Mix (Promega) and target specific primers (Supplementary Table 5).

### Detection of editing site by PCR and Sanger sequencing

K562 cDNA was used to amplify region of interest by PCR using FastStart Taq DNA polymerase (Roche) and gene specific primers (Supplementary Table 5). PCR cycle condition as follow: Initial denaturation at 95 °C for 5 minutes followed by 50 cycles of 95 °C for 30 seconds, 60 °C for 30 seconds and 72 °C for 1 minutes. PCR products were then purified by PCR purification kit (Qiagen) and subjected to Sanger sequencing (Axil sequencing). Presence of editing was visualized by sequencing chromatogram.

### Differential splicing analysis

We used the RNA-seq pipeline available at the CSI NGS Portal (42) to identify differential alternative splicing events between two samples. Splicing analysis was performed between ADAR1 KD and ADAR1 WT. The cutoff values (FDR q<0.05; |ΔSI|>10%) were used to identify differential splicing events and DAVID (43) was used for analyzing the enriched KEGG pathway. The overlapping background between ADAR1 and splicing events was based on the overlapping of ADAR1 peaks with all annotated splicing sites.

## Authors’ contributions

DGT, HY, CSW, HKT, and LYH conceived and designed the project. SJT, HKT and LC carried out the experiment. CSW, HY, and JL collected and analyzed the data. CSW, HY, HKT, LYH, WWT, QZ and LC contributed to the interpretation of the results. HKT, KTT and OA verified the analytical methods. QZ assisted with sample preparation. HY and DGT supervised the project. CSW and HY wrote the manuscript with support from LYH, KTT, SJT, OA, WWT and LC. All authors provided critical feedback and helped shape the research, analysis or manuscript.

## Ethics approval and consent to participate

Not applicable.

## Availability of data and materials

Following sequencing data sets were downloaded from ENCODE: (1) the ADAR1 K562 fRIP-seq data set and its control data set, (2) the strand-specific total RNA-seq from K562 cells, (3) ChIP-seq control from K562 cells, (4) total RNA-seq from ADAR1-depleted K562 cells. A list of data sets with ENCODE accession numbers was provided with Table S4 in Additional file 2.

## Additional files

### Additional file 1: supplementary figures

**Fig. S1.** Schematic flow for identification of ADAR1 RNA editing sites.

**Fig. S2.** Percentages of the overlapping number between A-to-I editing sites and ADAR1 peaks at different fold change cut-offs of fRIP-seq fold enrichment.

**Fig. S3.** Distribution of A-to-I editing sites located outside repeated regions.

**Fig. S4.** Profiling of differential splicing events in ADAR1 depleted cells.

**Fig. S5.** Validation of A-to-I sites at the downstream of ANLN in the ADAR1 RIP.

### Additional file 2: supplementary tables

**Table S1.** A-to-I editing sites in K562

**Table S2.** ADAR1 PIP-seq peaks in K562

**Table S3.** Gene associated ADAR1 peaks in IGR

**Table S4.** Datasets from public domains

**Table S5.** List of primers

## References

1. Wagner RW, Smith JE, Cooperman BS, Nishikura K. A double-stranded RNA unwinding activity introduces structural alterations by means of adenosine to inosine conversions in mammalian cells and Xenopus eggs. Proc Natl Acad Sci U S A. 1989;86(8):2647–51.

2. Ramaswami G, Li JB. RADAR: a rigorously annotated database of A-to-I RNA editing. Nucleic Acids Res. 2014;42(Database issue):D109–13.

3. Picardi E, D’Erchia AM, Lo Giudice C, Pesole G. REDIportal: a comprehensive database of A-to-I RNA editing events in humans. Nucleic Acids Res. 2017;45(D1):D750–D7.

4. Bazak L, Haviv A, Barak M, Jacob-Hirsch J, Deng P, Zhang R, et al. A-to-I RNA editing occurs at over a hundred million genomic sites, located in a majority of human genes. Genome Res. 2014;24(3):365–76.

5. Eisenberg E, Levanon EY. A-to-I RNA editing - immune protector and transcriptome diversifier. Nat Rev Genet. 2018;19(8):473–90.

6. Ramaswami G, Li JB. Identification of human RNA editing sites: A historical perspective. Methods. 2016;107:42–7.

7. Ramaswami G, Lin W, Piskol R, Tan MH, Davis C, Li JB. Accurate identification of human Alu and non-Alu RNA editing sites. Nat Methods. 2012;9(6):579–81.

8. Tan MH, Li Q, Shanmugam R, Piskol R, Kohler J, Young AN, et al. Dynamic landscape and regulation of RNA editing in mammals. Nature. 2017;550(7675):249–54.

9. Athanasiadis A, Rich A, Maas S. Widespread A-to-I RNA editing of Alu-containing mRNAs in the human transcriptome. PLoS Biol. 2004;2(12):e391.

10. Kim DD, Kim TT, Walsh T, Kobayashi Y, Matise TC, Buyske S, et al. Widespread RNA editing of embedded alu elements in the human transcriptome. Genome Res. 2004;14(9):1719–25.

11. Bahn JH, Ahn J, Lin X, Zhang Q, Lee JH, Civelek M, et al. Genomic analysis of ADAR1 binding and its involvement in multiple RNA processing pathways. Nat Commun. 2015;6:6355.

12. Nishikura K. A-to-I editing of coding and non-coding RNAs by ADARs. Nat Rev Mol Cell Biol. 2016;17(2):83–96.

13. Walkley CR, Li JB. Rewriting the transcriptome: adenosine-to-inosine RNA editing by ADARs. Genome Biol. 2017;18(1):205.

14. Liddicoat BJ, Piskol R, Chalk AM, Ramaswami G, Higuchi M, Hartner JC, et al. RNA editing by ADAR1 prevents MDA5 sensing of endogenous dsRNA as nonself. Science. 2015;349(6252):1115–20.

15. Rice GI, Kasher PR, Forte GM, Mannion NM, Greenwood SM, Szynkiewicz M, et al. Mutations in ADAR1 cause Aicardi-Goutieres syndrome associated with a type I interferon signature. Nat Genet. 2012;44(11):1243–8.

16. Bhate A, Sun T, Li JB. ADAR1: A New Target for Immuno-oncology Therapy. Mol Cell. 2019;73(5):866–8.

17. Lamers MM, van den Hoogen BG, Haagmans BL. ADAR1: “Editor-in-Chief” of Cytoplasmic Innate Immunity. Front Immunol. 2019;10:1763.

18. Ishizuka JJ, Manguso RT, Cheruiyot CK, Bi K, Panda A, Iracheta-Vellve A, et al. Loss of ADAR1 in tumours overcomes resistance to immune checkpoint blockade. Nature. 2019;565(7737):43–8.

19. Hsiao YE, Bahn JH, Yang Y, Lin X, Tran S, Yang EW, et al. RNA editing in nascent RNA affects pre-mRNA splicing. Genome Res. 2018;28(6):812–23.

20. D GH, Kelley DR, Tenen D, Bernstein B, Rinn JL. Widespread RNA binding by chromatin-associated proteins. Genome Biol. 2016;17:28.

21. Wissink EM, Vihervaara A, Tippens ND, Lis JT. Nascent RNA analyses: tracking transcription and its regulation. Nat Rev Genet. 2019.

22. Morgan JT, Fink GR, Bartel DP. Excised linear introns regulate growth in yeast. Nature. 2019;565(7741):606–11.

23. Kim U, Wang Y, Sanford T, Zeng Y, Nishikura K. Molecular cloning of cDNA for double-stranded RNA adenosine deaminase, a candidate enzyme for nuclear RNA editing. Proc Natl Acad Sci U S A. 1994;91(24):11457–61.

24. Patterson JB, Samuel CE. Expression and regulation by interferon of a double-stranded-RNA-specific adenosine deaminase from human cells: evidence for two forms of the deaminase. Mol Cell Biol. 1995;15(10):5376–88.

25. Daniel C, Veno MT, Ekdahl Y, Kjems J, Ohman M. A distant cis acting intronic element induces site-selective RNA editing. Nucleic Acids Res. 2012;40(19):9876–86.

26. Rueter SM, Dawson TR, Emeson RB. Regulation of alternative splicing by RNA editing. Nature. 1999;399(6731):75–80.

27. Licht K, Kapoor U, Mayrhofer E, Jantsch MF. Adenosine to Inosine editing frequency controlled by splicing efficiency. Nucleic Acids Res. 2016;44(13):6398–408.

28. Solomon O, Oren S, Safran M, Deshet-Unger N, Akiva P, Jacob-Hirsch J, et al. Global regulation of alternative splicing by adenosine deaminase acting on RNA (ADAR). RNA. 2013;19(5):591–604.

29. Rieder LE, Reenan RA. The intricate relationship between RNA structure, editing, and splicing. Semin Cell Dev Biol. 2012;23(3):281–8.

30. Bentley DL. Coupling mRNA processing with transcription in time and space. Nat Rev Genet. 2014;15(3):163–75.

31. Davis CA, Hitz BC, Sloan CA, Chan ET, Davidson JM, Gabdank I, et al. The Encyclopedia of DNA elements (ENCODE): data portal update. Nucleic Acids Res. 2018;46(D1):D794–D801.

32. Dobin A, Davis CA, Schlesinger F, Drenkow J, Zaleski C, Jha S, et al. STAR: ultrafast universal RNA-seq aligner. Bioinformatics. 2013;29(1):15–21.

33. Harrow J, Frankish A, Gonzalez JM, Tapanari E, Diekhans M, Kokocinski F, et al. GENCODE: the reference human genome annotation for The ENCODE Project. Genome Res. 2012;22(9):1760–74.

34. Li H, Handsaker B, Wysoker A, Fennell T, Ruan J, Homer N, et al. The Sequence Alignment/Map format and SAMtools. Bioinformatics. 2009;25(16):2078–9.

35. Quinlan AR, Hall IM. BEDTools: a flexible suite of utilities for comparing genomic features. Bioinformatics. 2010;26(6):841–2.

36. Picard toolkit. Broad Institute, GitHub repository. 2019.

37. Kent WJ, Zweig AS, Barber G, Hinrichs AS, Karolchik D. BigWig and BigBed: enabling browsing of large distributed datasets. Bioinformatics. 2010;26(17):2204–7.

38. Kent WJ, Sugnet CW, Furey TS, Roskin KM, Pringle TH, Zahler AM, et al. The human genome browser at UCSC. Genome Res. 2002;12(6):996–1006.

39. Raney BJ, Dreszer TR, Barber GP, Clawson H, Fujita PA, Wang T, et al. Track data hubs enable visualization of user-defined genome-wide annotations on the UCSC Genome Browser. Bioinformatics. 2014;30(7):1003–5.

40. Haeussler M, Zweig AS, Tyner C, Speir ML, Rosenbloom KR, Raney BJ, et al. The UCSC Genome Browser database: 2019 update. Nucleic Acids Res. 2019;47(D1):D853–D8.

41. Ramirez F, Ryan DP, Gruning B, Bhardwaj V, Kilpert F, Richter AS, et al. deepTools2: a next generation web server for deep-sequencing data analysis. Nucleic Acids Res. 2016;44(W1):W160–5.

42. An Ö, Tan K, Li Y, Li J, Wu CS, Zhang B, et al. CSI NGS Portal: An Online Platform for Automated NGS Data Analysis and Sharing. Preprints. 2019(2019100146).

43. Huang da W, Sherman BT, Lempicki RA. Systematic and integrative analysis of large gene lists using DAVID bioinformatics resources. Nat Protoc. 2009;4(1):44–57.

