## Supplementary figures for "Widespread interaction between ADAR1 and transcriptional byproducts"

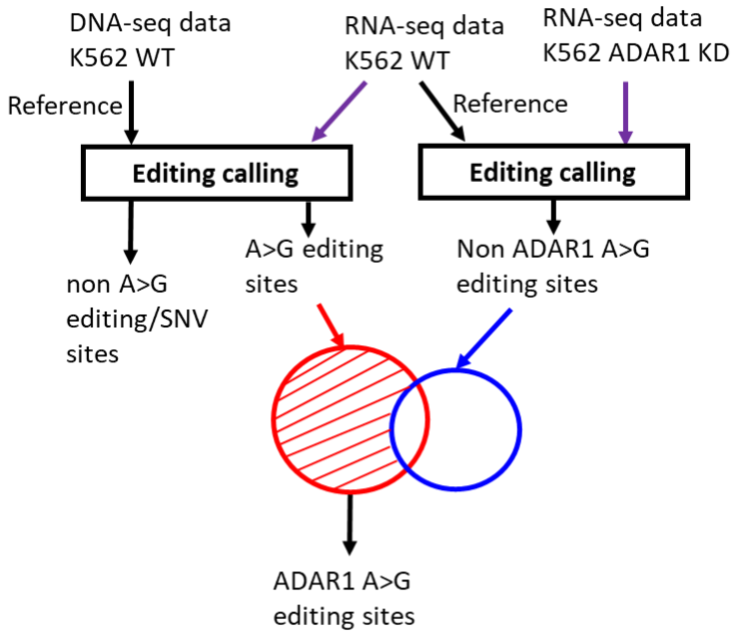


**Fig. S1.** Schematic flow for identification of ADAR1 RNA editing sites.

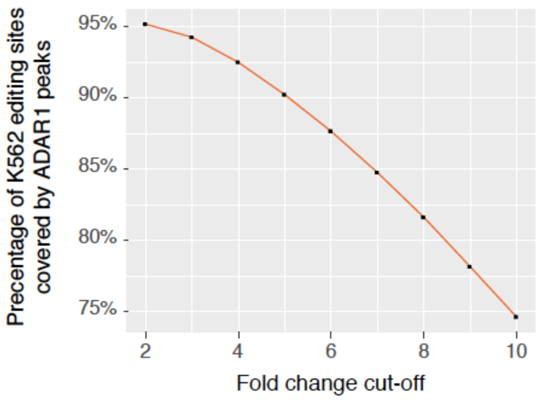


**Fig. S2.** Percentages of the overlapping number between A-to-I editing sites and ADAR1 peaks at different fold change cut-offs of fRIP-seq fold enrichment.

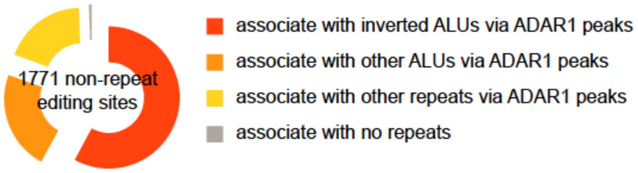


**Fig. S3.** Distribution of A-to-I editing sites located outside repeated regions.

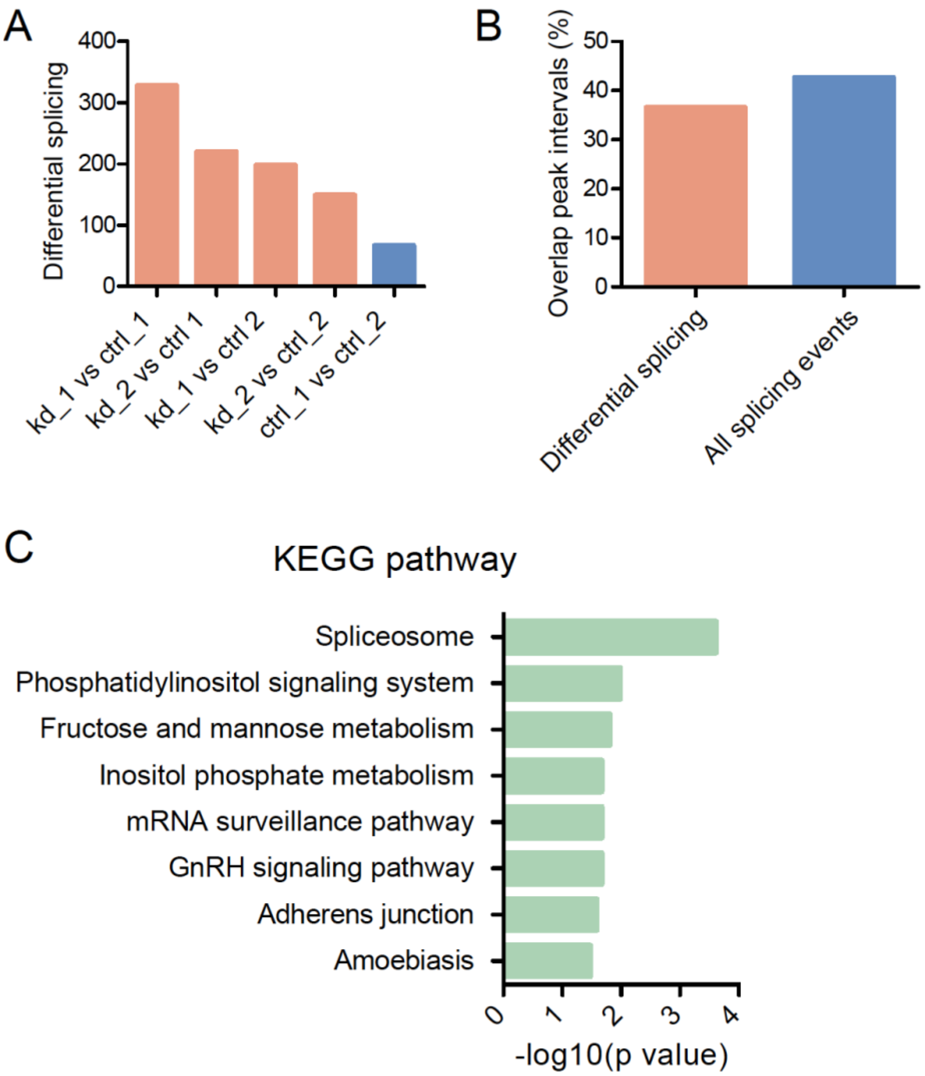


**Fig. S4.** Profiling of differential splicing events in ADAR1 depleted cells. (A) Number of differential splicing events (intron retention & exon skipping) between KD and WT. Differential splicing between control replicates was used as background noise. (B) No enriched overlapping of differential splicing events with ADAR1peaks compared to that of the background (all annotated splicing sites). (C) Enriched pathways in the GO analysis for all differentially spliced genes in ADAR1 KD versus control (WT).

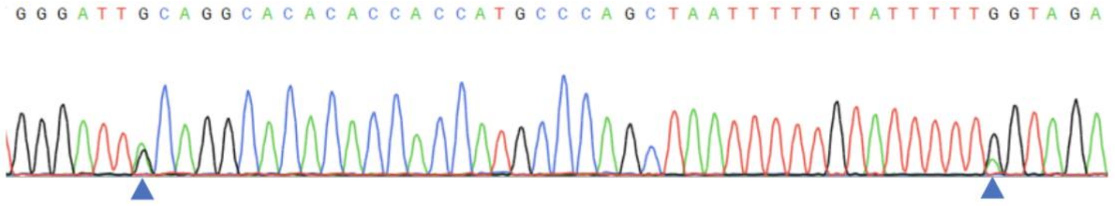


**Fig. S5.** Validation of A-to-I sites at the downstream of ANLN in the ADAR1 RIP.
